# RNA recognition by the E2 subunit of the chloroplast pyruvate dehydrogenase complex from Chlamydomonas

**DOI:** 10.1101/831339

**Authors:** Daniel Neusius, Laura Kleinknecht, Alexandra-Viola Bohne, Jörg Nickelsen

## Abstract

The dihydrolipoamide acetyltransferase subunit (DLA2) of the chloroplast puruvate dehydrogenase complex (cpPDC) from the green alga *Chlamydomonas reinhardtii* has previously been shown to possess a moonlighting activity in chloroplast gene expression. Exclusively under mixotrophic growth conditions, DLA2 forms part of an RNP particle with the *psbA* mRNA that encodes the D1 protein of the photosystem II reaction center. Here, we report on the further characterization of DLA2’s RNA-binding activity. Size-exclusion chromatography and Western analyses revealed that DLA2 is the only cpPDC subunit that shuttles between the metabolic cpPDC and the RNP complex. Microscale thermophoresis-based determination of RNA-binding affinities demonstrated that two DLA2 regions are crucial for RNA recognition, the peripheral E3-binding domain (E3BD) and the C-terminus of the catalytic domain. Specificity for the *psbA* RNA probe is conferred by the E3BD *in vitro*, as verified by competitive binding assays in the presence of an excess of the E3 (DLD2) of cpPDC. The data support a model in which an environmental trigger induces release of DLA2 from the cpPDC and its subsequent association with the *psbA* mRNA.

## Introduction

Cellular metabolism and gene expression are tightly interlinked. This becomes particularly important during acclimation processes required to cope with changes in the environment and/or the metabolic status of a cell. In mammals and bacteria, several lines of evidence indicate that gene expression is modulated by intermediates of carbon metabolism (1,2). In line with this, several glycolytic enzymes in the cytoplasm also act as regulators of transcription, mRNA stabilization and translation under specific conditions, a feature known as “moonlighting activity” (3). Since these enzymes often catalyze rate-limiting steps in metabolic pathways, they are ideally suited for mediating cross-talk between gene expression and C-metabolism under fluctuating environmental/metabolic conditions (4).

However, our knowledge of the relevant regulatory principles in chloroplasts – which serve as the major hub for C-metabolism in plants – is scarce (5). The few described examples of moonlighting factors in plastid gene expression include a sulphite reductase involved in the compaction of chloroplast nucleoids and in regulating their transcription (6). A second case is represented by the ribulose bisphosphate carboxylase/oxygenase (Rubisco), which catalyzes CO_2_ fixation in the first step in the Calvin cycle. Under oxidizing conditions, the large subunit of this enzymatic complex (RbcL) from the green alga *Chlamydomonas reinhardtii* acquires an RNA-binding activity that mediates auto-regulatory feedback control of *rbcL* mRNA translation and/or the formation of chloroplast stress granules, which sequester or degrade oxidized RNAs (7,8). Moreover, a plastid UMP kinase (PUMPKIN) has recently been described to bind to intron regions of chloroplast transcripts, thus linking RNA and pyrimidine metabolism in *Arabidopsis thaliana* (9).

A direct molecular connection between gene expression and fatty acid metabolism has been discovered in Chlamydomonas chloroplasts with the unexpected finding that the dihydrolipoyl acetyltransferase subunit (DLA2) of the chloroplast pyruvate dehydrogenase complex (cpPDC) possesses intrinsic RNA-binding activity (10). The cpPDC complex is a megadalton-sized complex consisting of multiple copies of three enzymatic components, i.e., the pyruvate dehydrogenase subunits E1α and E1β (PDC2 and PDH2 in *C. reinhardtii*), the dihydrolipoamide acetyltransferase E2 (DLA2 in *C. reinhardtii*), and a dihydrolipoyl dehydrogenase E3 (DLD2 in *C. reinhardtii*) (11,for an overview on PDC structures see 12). This giant enzyme complex catalyzes the oxidative decarboxylation of pyruvate to acetyl-CoA, the initial reaction in chloroplast fatty acid synthesis. However, under favorable mixotrophic growth conditions, i.e., in the presence of light as well as acetate, but not under photoautotrophic or heterotrophic conditions, a membrane-associated ribonucleoprotein (RNP) complex containing DLA2 was detected (10,13). This RNP is specifically formed with the 5’ untranslated region (UTR) of the *psbA* mRNA, which encodes the D1 protein of the photosystem II (PSII) reaction center (10,13). Moreover, it was shown that the *psbA* mRNA can inhibit cpPDC activity in chloroplast extracts, further supporting the idea of a reciprocal functional relationship between gene expression and fatty acid synthesis via a moonlighting activity of DLA2 (10). In line with this, DLA2 appears to be involved in an acetate-dependent relocalization of the *psbA* mRNA to a translation (T) zone in proximity to the chloroplast’s pyrenoid (10,14). Thus, it was hypothesized that elevated chloroplast acetyl-CoA levels signal the presence of adequate amounts of substrate for fatty acid synthesis by causing product inhibition of the cpPDC and its partial disassembly. This could make DLA2 available for RNP formation with the *psbA* mRNA, thus supporting its targeting to T-zones for PSII biogenesis. In this way, DLA2 would mediate the coordination of lipid and protein synthesis during the *de novo* generation of photosynthetic membranes in a light-and acetate-dependent manner (5,10).

Like other PDC E2 subunits, the DLA2 protein from *C. reinhardtii* has a multidomain structure, comprising an N-terminal lipoyl-binding domain, an E3 subunit-binding domain (E3BD), and a large C-terminal 2-oxo acid dehydrogenase catalytic domain of ~220 amino acids that forms the active site for acetyl-CoA production. Domains are connected by alanine-and proline-rich hinge regions which are approximately 50 and 70 residues in length (10,12). Using bioinformatical analyses, Bohne et al. (10) had also identified a previously unrecognized putative NAD^+^/RNA-binding motif (i.e., a Rossman-fold; (15) within the E3BD of DLA2 from *C. reinhardtii*, as well as in homologous proteins from other organisms.

The data presented here reveal that DLA2 shuttles between the cpPDC and the RNP complex. In this process, the E3 binding site of DLA2 plays a key role in the specific recognition of the chloroplast *psbA* mRNA, whereas the C-terminal region of the protein contributes significantly to its overall affinity for RNA.

## Results

### DLA2 is the only cpPDC subunit that forms an RNP complex

The DLA2 protein from *C. reinhardtii* was previously shown to be an active subunit of the cpPDC complex (10). However, when grown under mixotrophic conditions, DLA2 was additionally identified as a native component of an RNP that contains *psbA* mRNA (10,13).

To test whether other subunits of the cpPDC form part of this RNP, we determined the sizes of PDC2 (E1α)-, PDH2 (E1β)-and DLD2 (E3)-containing membrane-bound complexes by size-exclusion chromatography (SEC). As previously described, DLA2 is found in a membrane-associated, high-molecular-weight complex in the MDa range which includes an RNA moiety, as indicated by a shift in the size of the particle towards lower molecular-weight ranges upon treatment with RNases (Fig. 1; 10). In contrast to DLA2, the other cpPDC subunits form smaller complexes of ca. 700 kDa. Importantly, these complexes are insensitive to RNase digestion, indicating that they do not form RNPs (Fig.1). Therefore, DLA2 is most probably the only cpPDC subunit that binds to RNA and plays a role in the regulation of chloroplast gene expression.

**Figure 1.**
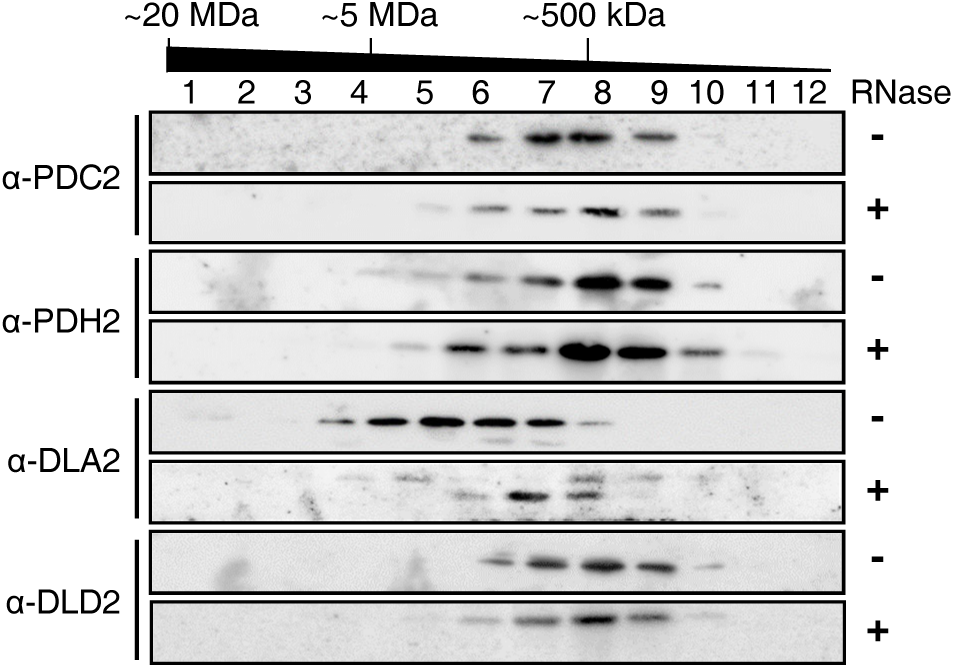
SEC analysis of cpPDC subunits. Thylakoids from the cell-wall-deficient wild-type strain CC-406 were solubilized and either treated with (+) RNase or not (-) before loading on the column. Fractions were collected and 1/10th of each fraction was loaded on an SDS gel. cpPDC subunits were detected by immunoblotting using antibodies indicated on the left.

The fact that DLA2 is the only cpPDC subunit engaged in growth condition-dependent RNP complex formation raises the question whether an additional pool of DLA2 is synthesized and/or the stoichiometry of cpPDC subunits changes when cells are growing mixotrophically. Alternatively, DLA2 could be released from the cpPDC under acetate-rich conditions in order to execute its moonlighting function in gene expression. To distinguish between these two possibilities, we quantified the steady-state levels of all cpPDC subunits by immunoblot analyses of total cell lysates of photo-, mixo-and heterotrophically grown wildtype Chlamydomonas cells. As shown in Fig. 2, levels of all subunits, including DLA2, remained constant, irrespective of the growth conditions employed. This indicates that there is no change in the size or allocation of the DLA2 pool under mixotrophic as compared to photoautotrophic or heterotrophic conditions. Hence, it appears most likely that mixotrophic conditions induce at least partial release of DLA2 from the metabolic enzyme complex, thereby enabling the protein to interact with the *psbA* mRNA.

**Figure 2.**
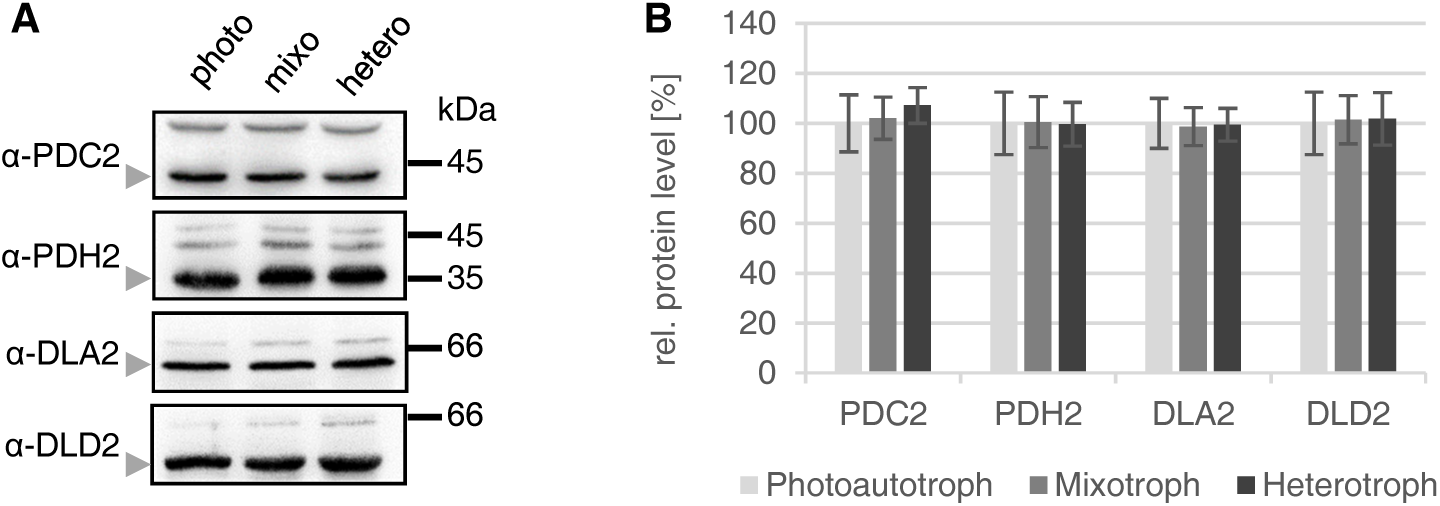
Quantification of cpPDC subunits under different growth conditions. **A.** Total proteins (15 μg) from wild-type cells grown under heterotrophic (hetero), mixotrophic (mixo) or photoautotrophic (photo) conditions were prepared as described before by Bohne et al. (10) fractionated by SDS-PAGE and subjected to immunoblot analysis. The indicated cpPDC subunits are marked by gray arrowheads. **B.** Quantification of immunoblot signals of three independent experiments. Standard deviations are shown and values for photoautotrophic growth were set to 100% for each protein.

### Mode of RNA binding to DLA2

To gain more detailed insights into how DLA2 binds to the *psbA* mRNA, microscale thermophoresis (MST) experiments were performed to quantify the affinity of various mutated versions of the protein for a Cy5-labelled *psbA* 5’ UTR RNA probe. As previously described, an A-rich region within the 5’ UTR of the *psbA* mRNA is essential for its recognition by DLA2 and D1 synthesis (13). Therefore, the probe used here covered 49 nucleotides of the *psbA* mRNA from positions −36 to +13 relative to the AUG start codon, which includes this A-stretch (compare Fig. 5A).

First, we determined the affinity of the wild-type DLA2 for this RNA fragment in comparison to that of an unrelated negative control protein, i.e., PratA, a cyanobacterial protein that lacks RNA-binding activity (16). For DLA2, MST traces showed a clear concentration-dependent fluorescence shift resulting in a typical saturation curve which is indicative of binding to the RNA probe (Fig. 3A+B). The KD value of 106 nM was slightly higher than that previously determined (51 nM) using filter-binding assays and a radioactive probe (Bohne et al., 2013). This might be due to the (5’ terminally) slightly shorter *psbA* probe used here and/or the different experimental setups (Fig. 4, DLA2; Bohne et al. 2013). In contrast, there was no concentration-dependent shift of MST traces for PratA (Fig. 3C+D), confirming that the effect seen for DLA2 is indeed caused by its interaction with the RNA probe.

**Figure 3.**
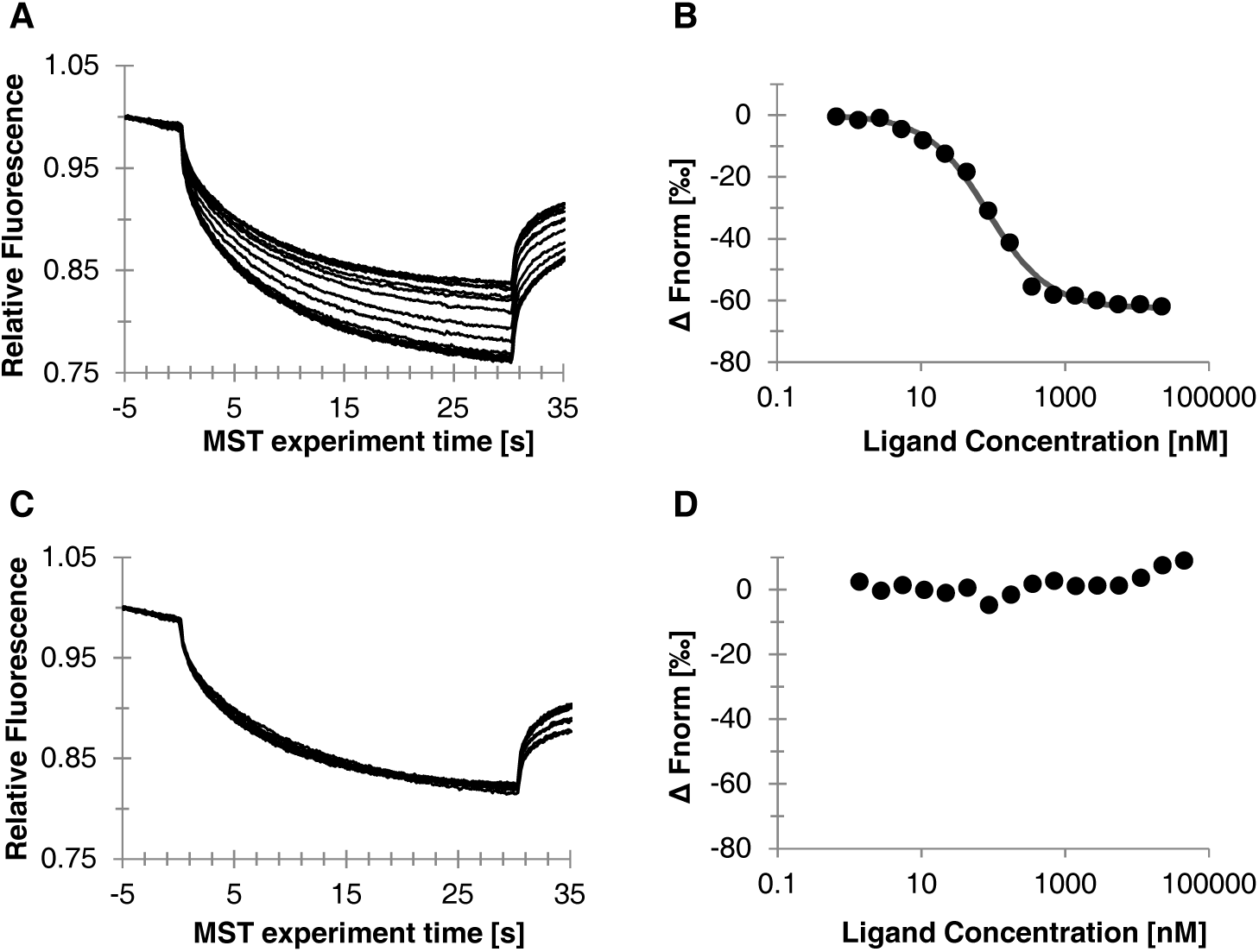
Microscale thermophoresis (MST) traces and RNA-binding curves of the recombinant DLA2 protein (A, B) and the *Synechocystis* protein PratA (16) as a negative control (C, D). **A, C.** MST traces are shown in black. **B, D.** The difference in fluorescence between the MST traces after 15 sec was evaluated and is depicted as black dots. The best fit generated by the NanoTemper software (NanoTemper, Munich) is depicted as a dark gray line.

**Figure 4.**
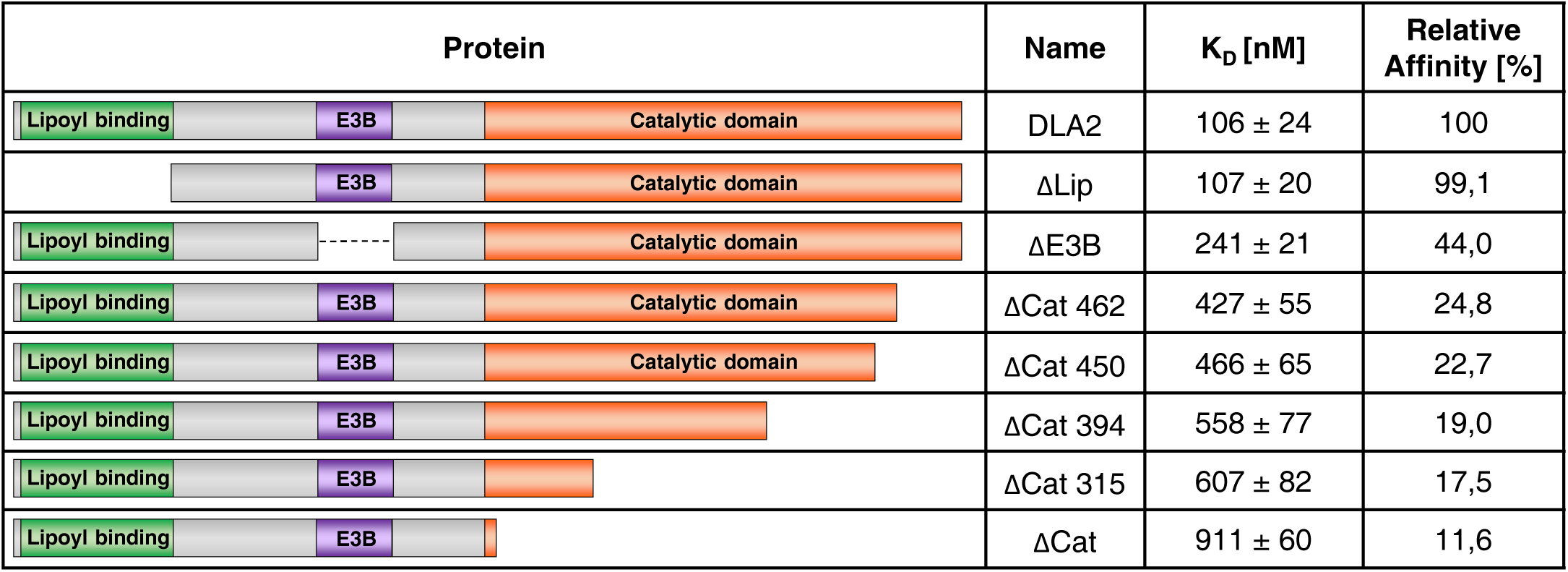
RNA-binding affinities of various mutant DLA2 proteins. The binding affinity of different DLA2 mutant proteins for a Cy5-labelled 5’UTR *psbA* probe was measured using microscale thermophoresis (MST). A schematic representation showing the domain structure of the analyzed recombinant proteins is displayed in the left column. Dissociation constant (K_D_) values are displayed as mean ± standard deviation of three independent experiments. For better comparability the binding affinity relative to the WT protein is shown. The amino acid numbers given in protein names refer to the first deleted amino acid in the respective protein. If no amino acid number is indicated the respective domain was deleted completely.

As mentioned above, DLA2 harbors three defined functional domains, i.e., the lipoyl-binding, E3B, and catalytic domains. We therefore tested mutated versions of DLA2 to identify the region(s) involved in RNA binding (Fig. 4; Supplemental Fig. S1). When the DLA2’s lipoyl-binding domain was deleted, no difference in RNA-binding affinity was detected relative to the wild-type protein; hence, this domain is apparently not involved in RNA binding (Fig. 4, ΔLip). Upon deletion of the E3BD, the RNA-binding affinity was severely diminished by 56% compared to the wild-type protein (Fig. 4, ΔE3B). Furthermore, when the catalytic domain was deleted, RNA binding was drastically decreased to 11,6% of the wild-type level (Fig. 4, ΔCat). These results indicate that both the E3BD and the catalytic domain contribute to RNA recognition.

To narrow down the RNA-binding region within the large catalytic domain of DLA2, we progressively deleted portions of it, starting from the C-terminus. Deletion of the last 33 amino acids alone led to a drastic reduction in RNA binding down to 24.8% of the wild-type level, indicating a crucial role of the C-terminal end for *psbA* mRNA recognition (Fig. 4; ΔCat 462). When the C-terminal region was further truncated (ΔCat 450, ΔCat 394, ΔCat 315, Fig 4), the affinity for RNA further decreased to 22,7%, 19,0% and 17,5% of the wild-type level, respectively (Fig. 4; ΔCat 450, ΔCat 394, ΔCat 315). This suggests that the whole catalytic domain, but especially the extreme C-terminal portion is required for the RNA-binding capacity of DLA2.

The existence of two apparently distinct RNA-binding regions in DLA2 raised the question whether both of them recognize the *psbA* mRNA probe with the same specificity. To test this, a mutated version of the *psbA* probe was synthesized in which the critical A-tract was replaced by a C-stretch (Fig. 5A). When this RNA probe was subjected to MST analysis, a significant reduction in binding affinity (to 42,7% of that measured for the native probe) was observed, in line with previous results (Fig. 5B; 13). A related, C-stretch-dependent reduction was also observed for the residual activity of the C-terminally truncated DLA2 version (Fig. 5B; ΔCat462). However, the DLA2 mutant lacking the E3BD was unable to discriminate between the two RNA probes, indicating that the E3BD actually confers specificity for *psbA* mRNA, whereas the C-terminus of DLA2 contributes to overall RNA binding (Fig. 5B; ΔE3BD).

**Figure 5.**
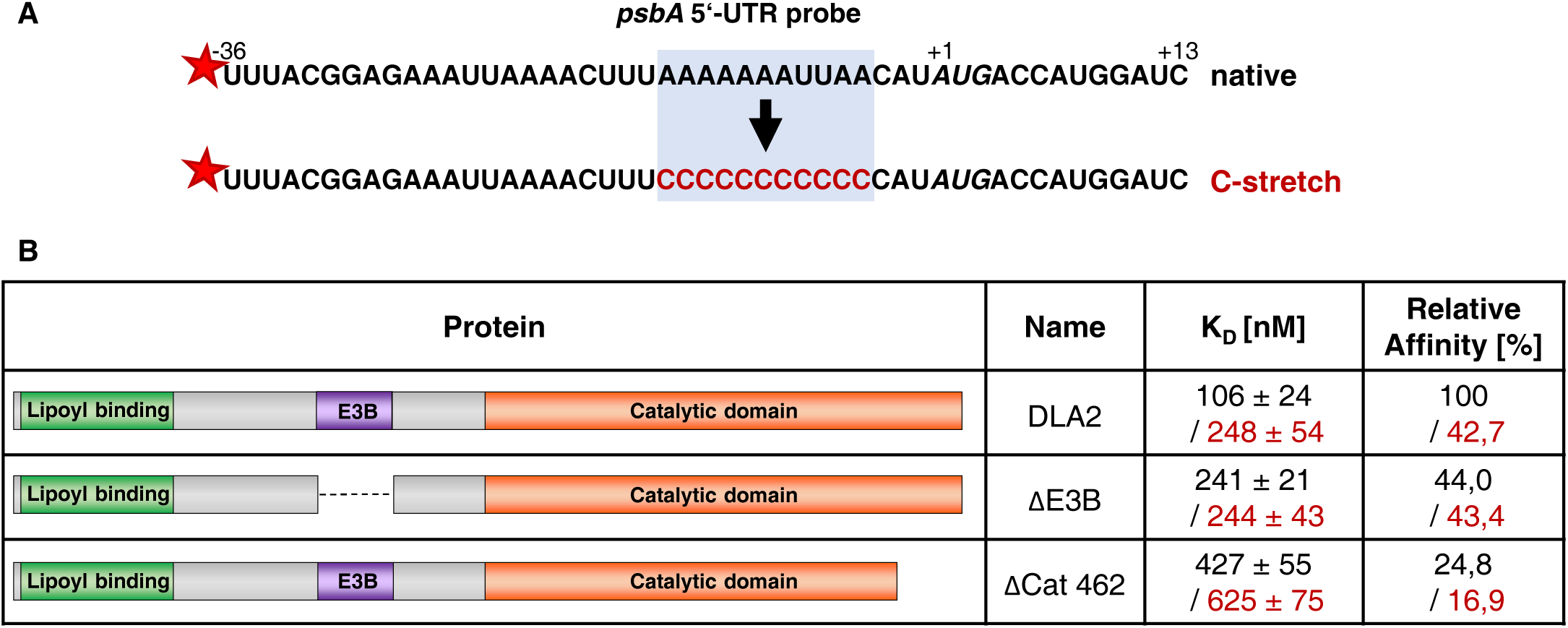
Specificity of RNA recognition by recombinant DLA2. **A.** Sequences of the native *psbA* 5’-UTR probe and the mutated probe with the C-stretch. Mutated nucleotides are depicted in red and positions relative to the start codon (italics) are marked above the sequence. **B.** Comparison of the binding affinities of different DLA2 mutant proteins for the native Cy5-labeled 5’UTR *psbA* probe and the mutated C-stretch probe. The means ± standard deviations (n=3) of the binding affinities to the native *psbA* probe are shown in black, while the values for the mutated C-stretch probe are shown in red. All affinity values indicated in the right column are relative to the binding affinity of the WT protein to the native *psbA* probe.

### *DLD2 and the* psbA *mRNA compete for binding to DLA2*

The RNA/protein interaction data strongly suggest that the *psbA* mRNA is recognized via DLA2’s C-terminus and the E3BD, which specifically recognizes the A-stretch essential for *psbA* translation (13,17). This scenario would predict that the *psbA* mRNA and the DLD2 (E3) subunit of the cpPDC compete for binding to the same DLA2 region (18). To test this hypothesis experimentally, we performed competitive RNA-binding experiments using recombinant DLA2 and DLD2, as well as the *psbA* 5’ UTR RNA probe. MST-based measurements of binding activities in such a three-component context proved difficult to interpret, so we turned to the UV crosslinking technique employed by Bohne et al., (2013) for the detection of DLA2 binding to radioactively labeled *psbA* RNA probe (Fig. 6). As previously reported, the apparent molecular weight of DLA2 in SDS gels is at 63 kDa, while its calculated mass is 47 kDa (10). Hence, the UV crosslinking signal of His-tagged DLA2 alone is detected in this size range (Fig. 6A, lane 3). The recombinant DLD2-GST fusion protein is in the similar size range, i.e., 63 kDa (see Experimental Procedures). However, no RNA binding signal was detected when DLD2 alone was assayed with the radioactive RNA probe, confirming the in-vivo observation that DLD2 is not found in any RNP (Fig. 1; Fig. 6A, lane 2). Nevertheless, when DLD2 was added in increasing amounts – up to tenfold molar excess – to the DLA2 RNA-binding reaction, the signal at 63 kDa was titrated out, indicating competition between DLD2 and *psbA* RNA for DLA2 (Fig. 6A, lanes 4-6). When the unrelated glutathione-S-transferase from *E. coli* was analyzed as a control protein, RNA binding of DLA2 was not affected, confirming that the observed competition effect is specific for DLD2 (Fig. 6B). Taken together, the data strongly support the idea that the DLD2 binding site on DLA2 represents the critical RNA interaction domain that confers DLA2’s moonlighting activity as a component of a specific RNP.

**Figure 6.**
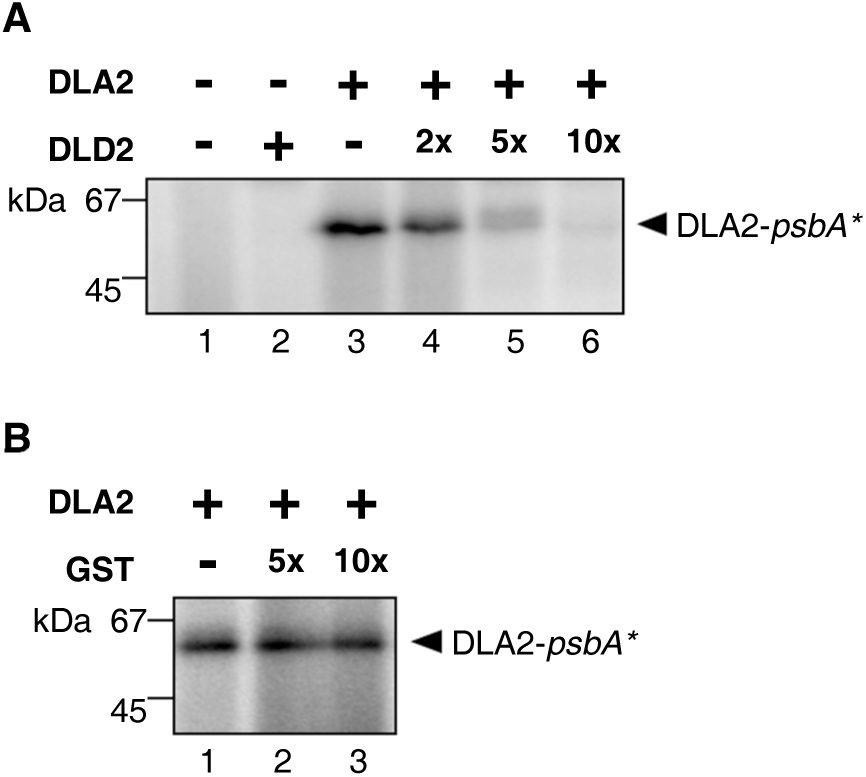
Quantification of competition for the *psbA* 5’ UTR RNA between recombinant DLD2 and DLA2. **A.** A radiolabeled *psbA* RNA probe was UV-crosslinked to 2 pmol DLA2 in the presence of 2, 5 and 10x molar excess of DLD2. To exclude binding of the *psbA* RNA probe to DLD2, a reaction using 100 ng of DLD2 instead of DLA2 was included as control. **B.** To show that the competing effect is caused specifically by DLD2, 5 and 10x molar excess of GST protein was added instead of DLD2 as negative control.

## Discussion

In recent years, the pyruvate dehydrogenase complex has attracted increasing attention as a moonlighting enzyme involved in the regulation of gene expression, in addition to its enzymatic role in acetyl-CoA production. For instance, subunits of human mitochondrial PDC have also been localized to the nucleus, where they appear to control gene transcription via interaction with transcription factors and/or histone acetylation (19,20). The cpPDC subunit DLA2 represents one example of a chloroplast C-metabolic enzyme that moonlights in plastid gene expression in the green alga *C. reinhardtii* (10). As proposed by Bohne et al. (10), thereby, DLA2 could mediate reciprocal molecular crosstalk between fatty acid synthesis and protein synthesis, enabling functional coordination of the two pathways for optimal thylakoid membrane biogenesis.

Here, we have shown that DLA2 is the only cpPDC subunit that forms an RNase-sensitive high-molecular-weight RNP complex in the MDa range, as assessed by SEC of solubilized chloroplast membranes (Fig. 1). Upon RNase digestion, DLA2 appears to comigrate with other subunits at lower molecular weights of ca. 700 kDa. This is, however, a smaller than the size expected for a fully assembled cpPDC, as its mitochondrial counterpart, as well as prokaryotic complexes, have been reported to exhibit sizes of several MDa (12). The detection of only smaller complexes of cpPDC subunits might be a feature of membrane-associated cpPDC units and/or might be attributable to the known instability of the cpPDC during its biochemical purification (10,21,22). However, the data revealed that the overall levels and, thus, the stoichiometry of all cpPDC subunits remain constant under different growth conditions (Fig. 2). This strongly supports the idea that DLA2 shuttles between cpPDC and RNP complexes, and argues against the existence of a separate DLA2 pool solely dedicated to RNP formation under mixotrophic growth conditions.

Previous bioinformatical predictions of potential RNA-binding domains within DLA2 using the RNAbindR tool identified two regions with high probability for RNA recognizing clusters of amino acids. These include the E3-binding domain spanning positions 188-224 (which contains a Rossmann fold) as well as the C-terminus between positions 450-457 (10; Supplemental Fig. S4). The measured binding affinities of several DLA2 versions determined in the present work confirm that both regions indeed play critical roles for RNA recognition. Deletion of E3BD reduced the affinity of DLA2 to less than 50% of the wildtype activity and, concomitantly, specificity for the important A-stretch within the *psbA* 5’UTR was completely lost (Fig. 5). Moreover, competition experiments verified that DLD2, the E3 subunit of the algal cpPDC, uses overlapping regions of DLA2 for its interaction when the metabolic enzyme is formed (Fig. 6).

Unexpectedly, removal of the last 33 amino acids from the C-terminus (up to position 462) is sufficient to cause a severe drop in RNA-binding activity (to 24,8% of the wild-type level) and has a larger effect than the deletion of the E3BD. Notably, the C-terminal truncation does not cover the residues (450-457) predicted to have the potential for RNA recognition. However, the truncation of the DLA2 region in the vicinity of positions 450-457 might already have a severe impact on their binding capacities. Alternatively, mutation of the C-terminus might affect the self-organization capacity of the catalytic domain, which is involved in the formation of the cubic E2 core of PDCs (11,12). If so, core formation would be required for RNA recognition by the E3BD, and the drastic drop in RNA binding by ΔCat 462 might be a more indirect effect. Nevertheless, deletions of increasingly larger segments from the DLA2 C-terminus resulted in a very gradual reduction in RNA binding, further supporting the idea of a role of the catalytic domain for interaction with the *psbA* RNA. Strikingly, the residual RNA-binding activity of the truncated ΔCat 462 version still conferred RNA specificity, as documented by the further reduction of the affinity to a mutant C-stretch RNA probe. Hence, the data suggest that the C-terminus has a decisive function for determining overall affinity to RNA, whereas the E3BD mediates specific recognition of the A-stretch within the *psbA* 5’ UTR.

In conclusion, the experimental findings support a model in which DLA2 is released from the metabolic cpPDC complex specifically under mixotrophic conditions, i.e., in the presence of both light and acetate, when cells are growing rapidly. To date, it is unclear what molecular signal triggers this initial step in DLA2’s transition to its moonlighting role in chloroplast gene expression. However, it appears likely that a weakening of the DLA2/DLD2 interaction is involved, which would give the *psbA* mRNA access to the E3BD region and may expose DLA2’s C-terminus. Clearly, more work is required to uncover the precise molecular events that enable DLA2 to serve as an RNA-binding protein as well as the precise ultrastructure of the DLA2/RNA complex.

### Experimental procedures

#### Growth of C. reinhardtii

Wild-type *C. reinhardtii* (strain CC-406) was grown in Tris/acetate/phosphate medium containing 1% sorbitol (TAPS) under mixotrophic or heterotrophic growth conditions (23). For photoautotrophic growth, cultures were cultivated in high salt minimal (HSM) medium (23). All liquid cultures were agitated under illumination at a fluence of 30 μmol photons m^−2^ s^−1^ at 23°C and harvested at a density of ca. 2 × 10^6^ cells/mL.

#### Antibody generation

To raise an antibody against the *C. reinhardtii* PDH2 protein, a fusion protein containing the glutathione-S-transferase (GST) was used as antigen. The expression vector was constructed as follows. A DNA fragment coding for amino acids 213-370 was amplified from cDNA by PCR using the primers PDH2 fw2 BamHI and PDH2 rev EcoRI. The resulting fragment was inserted into pGEX4T1 (GE Healthcare) via the restriction sites introduced by PCR. Overexpression was performed in *Escherichia coli* BL21 (DE3) cells at 27°C overnight, and the expressed protein was purified with Protino^®^ Glutathione Agarose 4B (Macherey-Nagel) according to the manufacturer’s instructions. The resulting protein fraction was used to immunize rabbits for the production of a polyclonal antiserum (Biogenes).

An antibody against the DLD2 protein was obtained essentially as described for PDH2. A DNA fragment coding for amino acids 409-585 was amplified using the primers DLD2 fw2 BamHI and DLD2 rev EcoRI. The fragment was inserted into pGEX4T1. The protein was overexpressed and purified as described above for PDH2 and used to immunize rabbits (Biogenes). For primer sequences and protein identifiers see Supplemental Table S1.

#### Chloroplast preparation and size exclusion chromatography (SEC)

Chloroplast preparation and subsequent SEC with or without RNase treatment was performed as described before (10). Solubilized thylakoid membranes were loaded through an online filter onto a Sephacryl S500 HR column (GE Healthcare), and elution was performed at 4°C with a buffer containing 50 mM KCl, 2.5 mM EDTA, 5 mM ε-aminocaproic acid, 0.1% Triton X-100, and 20 mM Tricine-KOH, pH 7.8, at a flow rate of 0.3 mL/min using an ÄKTApurifier 10 system (GE Healthcare). One tenth of each elution fraction was then subjected to immunoblotting.

#### Immunoblot analysis

Immunoblot analysis was performed using standard procedures. Protein concentrations were determined by the Bradford assay following the manufacturer’s (C. Roth) instructions.

PDC2 antibody was used in a 1/10,000 dilution, and the antibodies against PDH2 and DLD2 which were generated for this study were used in dilutions of 1/2000 and 1/5000, respectively. The DLA2 antibody was generated by Bohne et al. (10) and used in a 1/1000 dilution. The secondary antibody (anti-rabbit-HRP; #A9169, Sigma) was used in a dilution of 1/20,000. Quantification of immunoblot signals was done using the ImageQuant software (GE Healthcare).

#### Expression and purification of recombinant proteins

For the UV-crosslinking experiment, recombinant DLA2 fused to a His-tag was expressed as previously described by Bohne et al. (10) and purified with Protino^®^ Ni-NTA Agarose beads (Macherey-Nagel) according to the manufacturer’s instructions. For the expression of recombinant DLD2, a DNA fragment coding for the amino acids 241–585 was amplified using the primers DLD2 fw BamHI and DLD2 rev *EcoR*I (for primer sequences see Supplemental Table S1). Subsequently, the fragment was cloned into the pGEX-4T-1 plasmid using *BamH*I and *EcoR*I restriction sites and the resulting construct was transformed into *Escherichia coli* Rosetta cells (Novagen). DLD2 overexpression was performed at 18°C overnight after induction with 0.5 mM IPTG. GST-tagged proteins were purified with Protino^®^ Glutathione Agarose 4B. All constructs employed for the expression of the recombinant proteins used in MST were generated by the same procedure. Fragments coding for each amino-acid sequence were amplified by PCR using primers listed in Supplemental Table S1. To increase overexpression, a synthetic version of the DLA2 cDNA (Metabion International AG, Germany) that conformed to the codon usage of *Escherichia coli* was used as a template for PCRs. The PCR product was cloned into the vector pET-28b-Sumo (24) using *BamH*I and *Sal*I restriction sites, and the resulting construct was transformed into *Escherichia coli* BL21 (DE3) cells. For the pSUMO: DLA2 ΔE3B construct expressing the ΔE3B protein, the 5’ region of the cDNA was amplified using the primers DLA2 cDNA fw 2 BamHI and DLA2 - E3B rev SalI and cloned into pET-28b-Sumo. The 3’ region was amplified using the primers DLA2-E3B fw SalI and DLA2 cDNA rev SalI and cloned downstream of the 5’ region via the restriction site *Sal*I. Overexpression of all recombinant proteins used for MST was performed at 12°C overnight after induction with 1 mM IPTG. His-Sumo-tagged proteins were purified with Protino^®^ Ni-NTA agarose beads, followed by the removal of the His-Sumo tag by proteolytic digestion with Sumo protease at 4°C overnight. All primer sequences used for the generation of the constructs as well as amino acid coverage of resulting proteins are listed in Supplemental Table S1.

#### Microscale thermophoresis

Microscale thermophoresis (MST) was used to quantify the binding affinities of Cy5-labeled *psbA* 5’UTR probes for wild-type and mutant DLA2 proteins. Synthetic RNAs were purchased from Metabion International AG. To guarantee high comparability between different samples, all recombinant proteins were overexpressed, purified and measured at the same time. The concentration of purified recombinant proteins was adjusted to 44 μM using MST buffer (50 mM Tris/HCl pH 7.8, 60 mM KCl, 10 mM MgCl_2_, 0.05% Tween 20). Subsequently the fluorescently labeled RNA probe was diluted to 20 nM in MST buffer and decreasing amounts of the respective proteins were incubated with the RNA. Afterwards samples were loaded into MST NT.115 premium glass capillaries (NanoTemper) and ran at 40% LED power and 20% MST power at RT in a Monolith NT.115 instrument (NanoTemper) at the Bioanalytic Service Unit of the Biocenter of the Ludwig-Maximilians-Universität, Munich. KD values were calculated using the MO. Affinity Analysis software (NanoTemper).

#### In Vitro Synthesis of RNA and UV CrossLinking

*In vitro* synthesis of RNA and UV cross-linking experiments were basically performed as described by Zerges and Rochaix (25) and Bohne et al. (10). The DNA template for the *in vitro* synthesis of the *psbA* 5’ UTR RNA probe was generated by PCR using the primers: T7psbA5 and 2054-psbA (for sequences see Supplemental Table S1. Binding reactions (20 μl) were performed at RT for 10 min and contained 20 mM HEPES/KOH, pH 7.8, 5 mM MgCl_2_, 60 mM KCl, and 100 ng of protein unless indicated otherwise. Each reaction contained 100 kcpm of ^32^P-RNA probe. For competition experiments, the indicated amounts of competitor protein were added to the reaction. Radiolabeled RNA and DLA2 were mixed prior to the addition of competing proteins.

## Acknowledgements

The authors thank P. Nixon for kindly providing PDC2 antiserum as well as Paul Irmer and Metabion International AG for providing codon-adapted DLA2 cDNA fragments.

## Author contributions

JN and AVB designed the experiments. DN and LK performed the experiments and analyzed the results. JN, AVB and DN wrote the paper.

## Footnotes

The nucleotide sequence for the synthetic version of the DLA2 cDNA codon-adapted to *Escherichia coli* has been deposited in the GenBank database under GenBank Accession Number MN642093.

Financial support from the Deutsche Forschungsgemeinschaft to J.N. (TRR175-A06) is acknowledged. L.K. was supported by a fellowship from the Deutsche Studienstiftung. The authors declare that they have no conflicts of interest in relation to the contents of this article.

The abbreviations used are: cpPDC, chloroplast pyruvate dehydrogenase complex; DLA, dihydrolipoamide acetyltransferase; GST, glutathione-S-transferase; HMW, high molecular weight; HSM, high-salt minimal; MST, microscale thermophoresis; PDH, pyruvate dehydrogenase; PSII, photosystem II; RBP, RNA-binding protein; RNP, ribonucleoprotein; SEC, size exclusion chromatography; K_D_, dissociation constant

## The abbreviations

cpPDC: chloroplast pyruvate dehydrogenase complex
DLA: dihydrolipoamide acetyltransferase
GST: glutathione-S-transferase
HMW: high molecular weight
HSM: high-salt minimal
MST: microscale thermophoresis
PDH: pyruvate dehydrogenase
PSII: photosystem II
RBP: RNA-binding protein
RNP: ribonucleoprotein
SEC: size exclusion chromatography

## Supporting Information

**Supplemental Table S1**. Primers used in this study.

**Supplemental Figure S1**. Coomassie brilliant blue-stained SDS gels demonstrating the purification of recombinant proteins used for MST analysis.

**Supplemental Figure S2**. MST-binding curves resulting from interaction of the mutant versions of DLA2 with the native *psbA* 5’-UTR probe.

**Supplemental Figure S3**. MST-binding curves resulting from interaction of different DLA2 versions with the mutated C-stretch *psbA* probe.

**Supplemental Figure S4**. Prediction of RNA-binding residues in the DLA2 protein from *C. reinhardtii* (adapted from Bohne et al. (10)).

