## Supporting Information for "RNA recognition by the E2 subunit of the chloroplast pyruvate dehydrogenase complex from Chlamydomonas"

**Supplemental Table S1. Primers used in this study.**

**Supplemental Figure S1. Coomassie blue stained SDS-Pages of the elution fraction of each recombinant protein used for MST analysis.**

**Supplemental Figure S2. Binding curves of the DLA2 mutant versions to the native *psbA* 5'-UTR probe.**

**Supplemental Figure S3. Binding curves of different DLA2 versions to the mutated C-stretch *psbA* probe.**

**Supplemental Figure S4. Prediction of RNA binding residues in the DLA2 protein from *C. reinhardtii (*adapted from Bohne et al. (1)*).***

**Supplemental Table S1. Primers used in this study.**

| Construct | Primer name and sequence (5'→ 3') | Protein | Amino acid coverage |
|---|---|---|---|
| pGEX: PDH2 Antigen | PDH2 fw2 BamHI (AA GGA TCC GTT CTG CTG TAC AAC GTG AAG G)<br>PDH2 rev EcoRI (AAGAATTCTGCTTGTGCTGCTGCTGCTTCG) | PDH2 Antigen | Aa 213-370[a] |
| pGEX: DLD2 Antigen | DLD2 fw2 BamHI (AA GGA TCC GAC GCC AAC GGC AAG TAC ATG CT)<br>DLD2 rev EcoRI (AAGAATTCGCGGCGGCCACCGCCGGAGG) | DLD2 Antigen | Aa 409-585 [b] |
| pGEX: DLD2 241-585 | DLD2 fw BamHI (AAGGATCCGACGGCAAGACCGTGTTCACCTC)<br>DLD2 rev EcoRI (AAGAATTCGCGGCGGCCACCGCCGGAGG) | DLD2 | Aa 241-585[b] |
| pSUMO:DLA2 WT | DLA2 cDNA fw 2 BamHI (TGGTGGATCCAATGCGGTTAAAGA)<br>DLA2 cDNA rev SalI (CTCGAGGCGCGTCGACGAATTCTTA) | DLA2 | Aa 32-494[c] |
| pSUMO: DLA2 ΔLip | DLA2-Lip fw BamHI (GGATCCGAAGAGGCGAAAAAGAAAGC)<br>DLA2 cDNA rev SalI (CTCGAGGCGCGTCGACGAATTCTTA) | ΔLip | Aa 116-494[c] |
| pSUMO: DLA2 ΔE3B | DLA2 cDNA fw 2 BamHI (TGGTGGATCCAATGCGGTTAAAGA)<br>DLA2-E3B rev SalI (GTCGACCCCGTCTGCACGGCCAAC) | ΔE3B | Aa 32-184[c] |
| pSUMO: DLA2 ΔE3B | DLA2-E3B fw SalI (5'-GTCGACGCAGCAGGCAAGGCTGTT)<br>DLA2 cDNA rev SalI (5'-CTCGAGGCGCGTCGACGAATTCTTA) | ΔE3B | Aa 224-494[c] |
| pSUMO: DLA2 ΔCat Aa461 | DLA2 cDNA fw 2 BamHI (TGGTGGATCCAATGCGGTTAAAGA)<br>DLA2 Cat Domain rev4 SalI (GCTGTCGACTTAGACGTTCATAACCTTCTTAACAC) | ΔCat Aa461 | Aa 32-461[c] |
| pSUMO: DLA2 ΔCat Aa449 | DLA2 cDNA fw 2 BamHI (TGGTGGATCCAATGCGGTTAAAGA)<br>DLA2 Cat Domain rev3 SalI (GCTGTCGACTTACGGAGACGCCACAACGGT) | ΔCat Aa449 | Aa 32-449[c] |
| pSUMO: DLA2 ΔCat Aa393 | DLA2 cDNA fw 2 BamHI (TGGTGGATCCAATGCGGTTAAAGA)<br>DLA2 Cat Domain rev2 SalI (GCTGTCGACTTACACCAGGTCGGCCCAGTT) | ΔCat Aa393 | Aa 32-393[c] |
| pSUMO: DLA2 ΔCat Aa314 | DLA2 cDNA fw 2 BamHI (TGGTGGATCCAATGCGGTTAAAGA)<br>DLA2 Cat Domain rev1 SalI (GCTGTCGACTTACAGCTGCTGGTACAGCGCA) | ΔCat Aa314 | Aa 32-314[c] |
| pSUMO: DLA2 ΔCat Domain | DLA2 cDNA fw 2 BamHI (TGGTGGATCCAATGCGGTTAAAGA)<br>DLA2-Cat rev SalI (GTCGACCTCGAGTTAGAGTTCGGAGACCGTGGTA) | ΔCat | Aa 32-269[c] |
| pSUMO: PratA | PratA-oe-fw BamHI (AAGGATCCAATCTTCCTGACGTTACC)<br>PratA-oe-rev PstI (AACTGCAGTTAGAGATTATCCAGCTTTTCT) | PratA | Aa 39-383[d] |
| *psbA* 5' UTR RNA probe (UV-crosslinking) | T7psbA5 (gtaatacgactcactatagggTACCATGCTTTTAATAGAAG)<br>2054-psbA (GATCCATGGTCATATGTTAATTTTTTTAAAG) | | |

Restriction sites introduced for cloning are underlined. Amino acid coverage given is related to the *C. reinhardtii* protein sequences obtained from Phytozome (v5.5): [a]PDH2, Cre03.g194200; [b]DLD2, Cre01.g016514; [c]DLA2, Cre03.g158900. The *Synechocystis* sp. PCC 6803 protein sequence of [d]PratA (*slr2048)* was obtained from cyanobase.

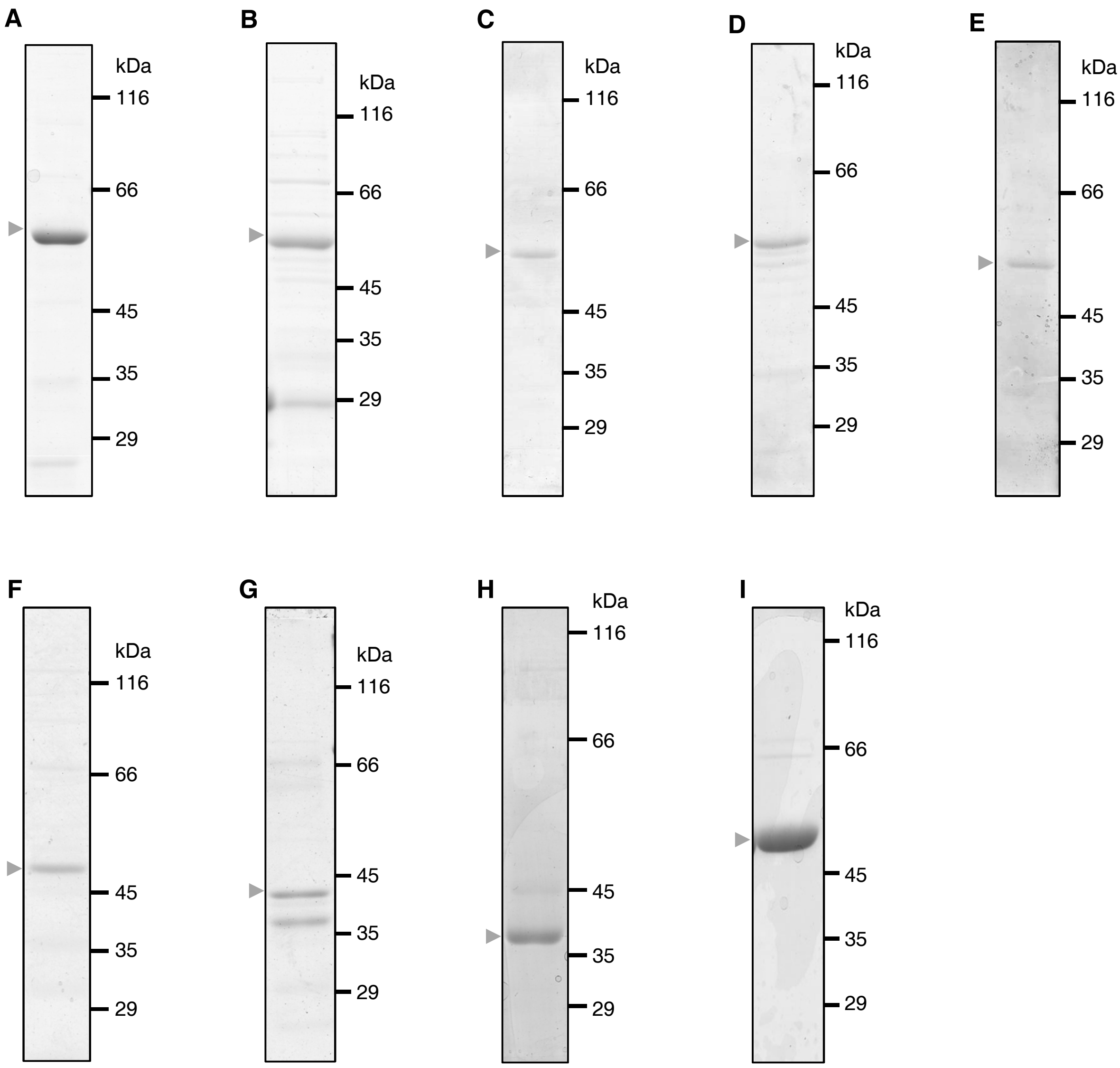


**Supplemental Figure S1. Coomassie brilliant blue-stained SDS gels demonstrating the purification of recombinant proteins used for MST analysis.** Elution fractions of recombinant wild-type DLA2 (A), ΔLip (B), ΔE3B (C), ΔCat 315 (D), ΔCat 394 (E), ΔCat 450 (F), ΔCat 462 (G), ΔCat (H) and PratA (I) were separated by SDS-PAGE and subsequently stained with Coomassie brilliant blue R-250. Samples were run alongside a molecular mass marker indicated in kDa.

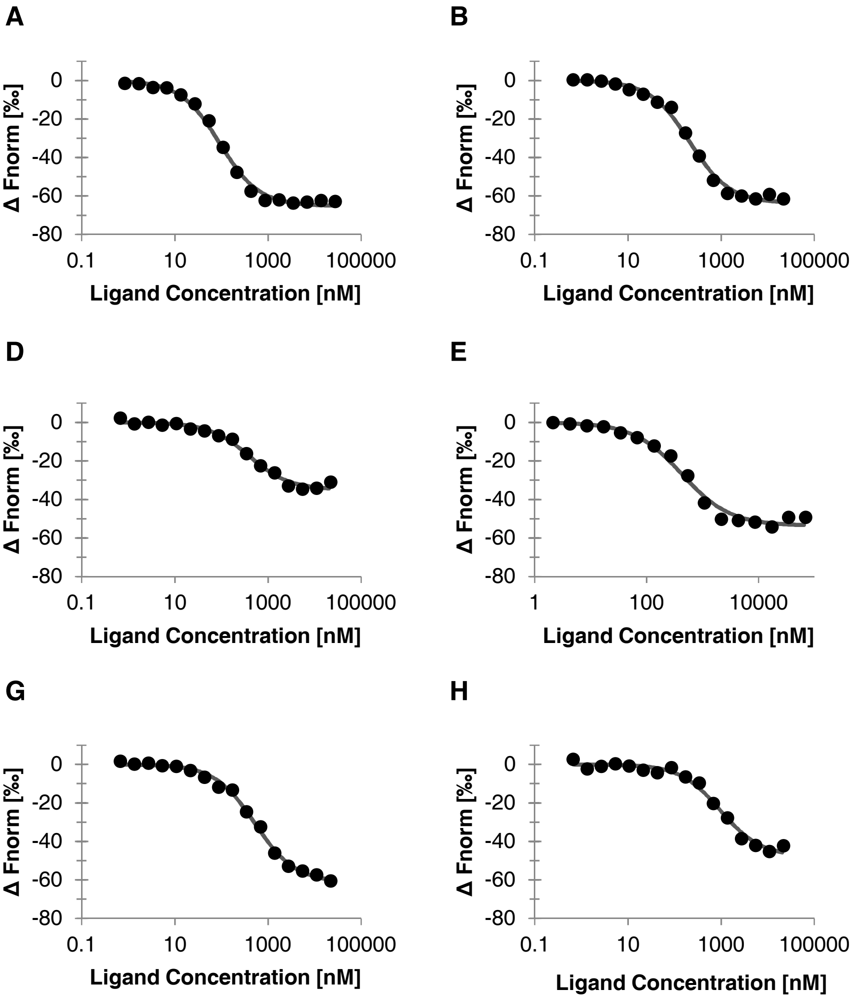


**Supplemental Figure S2. MST-binding curves resulting from interaction of the mutant versions of DLA2 with the native *psbA* 5'-UTR probe.** Shown are the binding curves for the ΔLip (A), ΔE3B (B), ΔLip&E3B (C), ΔCat 462 (D), ΔCat 450 (E), ΔCat 394 (F), ΔCat 315 (G) and the ΔCat (H) protein to a Cy5-labeled RNA probe. Measurement points of the fluorescence difference after 15 sec are depicted as black dots and the „Fit" generated by the NanoTemper software is displayed as a dark grey line. Curves are representative for 3 independent experiments.

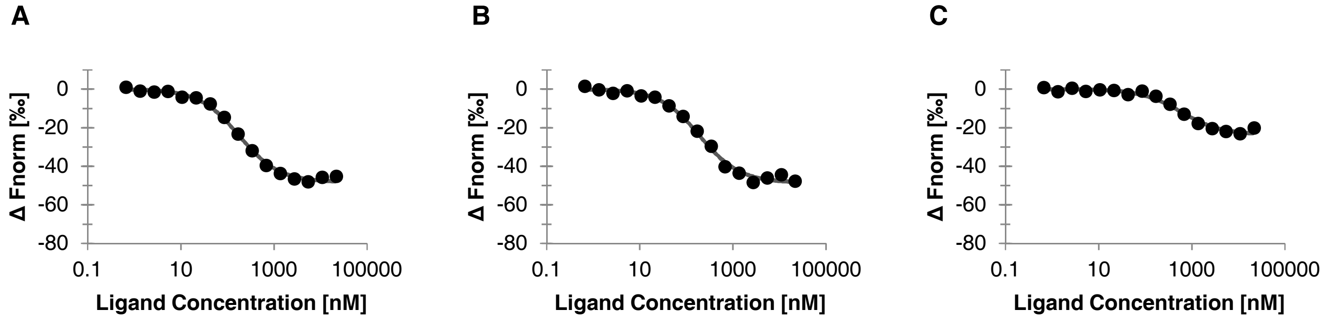


**Supplemental Figure S3. MST-binding curves resulting from interaction of different DLA2 versions with the mutated C-stretch *psbA* probe.** Binding curves for the wild-type DLA2 (A), ΔE3B (B), and ΔCat 462 (C) protein to the mutated Cy5-labeled RNA probe. Measurement points of the fluorescence difference after 15 sec are depicted as black dots and the „Fit“ generated by the NanoTemper software is displayed as a dark grey line. Curves are representative for 3 independent experiments.

```
         Transit peptide  20                   40  Lipoyl attachement  60
MQATTRVPAK SGVSSSAKRV AASGRRVLVV PNAVKDVFMP ALSSTMTEGK IVSWLKNVGD
-+--++++++ +++++++++- +++-+----- ---------- ---------- ----------

                    80                   100                   120
KVKKGEALVV VESDKADMDV ESFADGILGA IVVQEGERAV VGAPIAFVAE NANEVEEAKK
---------- ---------- ---------- ---------- ---------- ----------

                    140                  160                   180
KAAAAGGAAP AAAPAAAAAP APAPAAAPTP APAAAPAPAP APVAAPAPPA PTPVPAAPVG
---------- ---------- ---------- ---------- ---------- ----------

                    200  E3 binding region  220                 240
RADGRIVATP YAKQLAKDLK VDLATVAGTG PNGRITAADV EARAAGKAVA PAAAPAAAPA
---------- +-++--+--+ -------+++ ++++++---- --+---+--- ----------

                    260                  280                   300
AVAAAPAAAP AKAAAPAAAP AKATTVSELR GTTKPFSTLQ AAVARNMNES LKVPEFRVSY
---------- ---------- ---------- ---------- ---------- ----------

                    320  Catalytic domain  340                   360
AITTDKLDAL YQQLKPKGVT MTALLAKACG VALAKHPLLY AACTPDGNGI TYSSQINVAL
---------- ---------- ---------- ---------- ---------- ----------

                    380                  400                   420
AVAMPDGGLI TPVLKNADST DLYQMSRNWA DLVKRARSKQ LQPDEYNSGN FTISNLGMYG
---------- ---------- ---------- ---------- ---------- ----------

                    440                  460                   480
VETFDAILPP GTAAIMAVGG SKPTVVASPD GMIGVKKVMN VNLTADHRIV YGADAAEFLQ
---------- ---------- ---------+ +----++--- ---------- ----------

TLKAVIENPD QLLF
---------- ----
```

**Supplemental Figure S4. Prediction of RNA-binding residues in the DLA2 protein from *C. reinhardtii* (adapted from Bohne et al. (10)).** Possible RNA binding residues were previously predicted by using the RNAbindR tool (http://ailab-projects1.ist.psu.edu:8080/FastRNABindR/). Plus signs below the amino acid sequences indicate the predicted RNA-binding residues, whereas minus signs indicate a low probability of RNA binding. Plus signs within the processed protein (without transit peptide) were additionally highlighted in yellow. The conserved domains for lipoyl attachment, E3 binding, and the 2-oxo acid dehydrogenase catalytic domain are marked above the sequence by green, purple, and orange bars, respectively. The transit peptide is marked in blue.
